# Styx: A multi-language API Generator for Command-Line Tools

**DOI:** 10.1101/2025.07.24.666435

**Authors:** Florian Rupprecht, Jason Kai, Biraj Shrestha, Steven Giavasis, Ting Xu, Tristan Glatard, Michael P Milham, Gregory Kiar

## Abstract

In numerous scientific domains, established tools have often been developed with complex command-line interfaces. Such is the case for brain imaging and bioinformatics, making the use of powerful legacy tools in modern workflow paradigms challenging. We present (i) Styx, a compiler for generating language-native wrapper functions from static tool metadata, leading to seamless integration of command-line tools within the data science ecosystem. Alongside Styx, we have created (ii) NiWrap, a collection of more than 1900 neuroimaging command-line function descriptions as a proof-of-concept implementation. These interfaces, available in Python, R, and TypeScript (available at https://github.com/styx-api), significantly reduce the complexity of writing and interpreting software pipelines, particularly when composing workflows across packages with distinct API standards. The compiler architecture of Styx facilitates maintainability and portability across computing environments. As with all metadata-dependent infrastructure, creating sufficient metadata annotations remains a barrier to adoption. Accordingly, NiWrap demonstrates approaches that lower this barrier through direct source code extraction and LLM-assisted documentation parsing. Together, Styx and NiWrap offer a sustainable solution for interfacing diverse command-line tools with modern data science ecosystems. This modular approach enhances reproducibility and efficiency in pipeline development while ensuring portability across computing environments and programming languages.

## Introduction

Command-line interfaces (CLIs) have been the predominant distribution mechanism for scientific computing tools across numerous domains, particularly in neuroimaging and bioinformatics. These tools, ranging from established packages like FSL and AFNI to precisely focused software like ANTs and MRTrix3, collectively represent decades of algorithmic development and validation. However, integrating these diverse command-line tools into modern data science workflows presents significant challenges.

The primary challenge stems from the fundamental mismatch between CLI design and contemporary programming practices. While Python and R have become the de facto standards for developing data processing pipelines (Harris et al., 2020; McKinney, 2010; Pedregosa et al., 2011; Wickham et al., 2019), command-line tools require manual construction of arguments, file-based input/output operations, and domain-specific knowledge of each tool’s unique interface conventions. This gap creates barriers to reproducibility, increases development time, and raises the potential for errors in complex analytical pipelines.

In a comprehensive pipeline such as C-PAC (Craddock et al., 2013), approximately 100 unique command line interfaces across multiple software packages are integrated. Even this complex pipeline scratches the surface of tools used in neuroimaging, which Table 1 summarizes across several major packages. Manually creating and maintaining wrapper code for each tool represents a substantial burden that grows with each software update.

**Table 1.** Number of command-line interfaces in common neuroimaging software packages.

| Software package | Total CLIs |
| --- | --- |
| AFNI (Cox, 1996) | 611 |
| FreeSurfer (Fischl, 2012) | 800 |
| FSL (Jenkinson et al., 2012) | 310 |
| Connectome Workbench (Marcus et al., 2013) | 202 |
| MRTrix3 (Tournier et al., 2019) | 121 |
| ANTs (Avants et al., 2009) | 113 |
| Other tools* | 12 |

\* Includes Convert3D (Yushkevich et al., 2006) (2), NiftyReg (Modat et al., 2010) (7), Greedy (Yushkevich et al., 2006) (1), MRTrix3Tissue (Dhollander et al., 2021) (1), and dcm2niix (Li et al., 2016) (1)

Several approaches have emerged to address these integration challenges, each providing partial solutions. Nipype (Gorgolewski et al., 2011) pioneered the concept of Python wrappers for neuroimaging tools, offering both a workflow engine and hand-written interfaces. However, these interfaces require manual creation and maintenance for each tool, are limited to Python, and must be updated with each software release. The ongoing transition to Pydra (Jarecka et al., 2024) as Nipype’s successor further highlights the maintenance challenges of this approach.

Snakemake (Köster and Rahmann, 2018) takes a different approach, embedding shell commands within a domain-specific workflow language. While this reduces the need for explicit wrappers, it requires users to understand both the workflow language and the underlying command-line syntax, and tool invocations must be reimplemented in each workflow.

At a more fundamental level, Boutiques (Glatard et al., 2018) addresses the lack of standardized tool descriptions by providing a JSON-based specification for command-line interfaces. This machine-readable format enables tool interoperability and validation but still requires manual creation of descriptions and doesn’t directly provide programming language bindings.

We present Styx and NiWrap as a comprehensive solution that builds upon these existing approaches while addressing their limitations. Styx is a compiler that automatically generates native programming language interfaces from standardized tool descriptions, currently supporting Python, R, and TypeScript. NiWrap demonstrates this approach at scale with over 1,900 neuroimaging tool descriptions. Together, they enable seamless integration of command-line tools into modern data science workflows while maintaining portability across languages and computing environments. To contextualize our approach, we first examine existing tool description standards and their limitations in supporting multi-language wrapper generation.

## Tool Description Standards

For this manuscript, we focus on frameworks with explicit support for command-line utilities core to the neuroimaging and bioinformatics communities: Nipype and Snakemake.

Nipype (Gorgolewski et al., 2011; Esteban et al., 2022) is a Python framework that features both a Directed Acyclic Graph (DAG)-based dataflow engine and a collection of hand-written wrappers for neuroimaging tools. While using Nipype’s wrappers independently of its workflow engine is possible, it incurs overhead difficult to justify unless operating within a DAG-based workflow context. To address this, the team has begun to decouple the dataflow engine from the wrapper collection; the Pydra workflow engine (Jarecka et al., 2024) began development in 2018 as Nipype’s successor. However, manually porting all Nipype Interfaces to Pydra poses significant challenges requiring substantial maintenance efforts.

Snakemake (Köster and Rahmann, 2018), used mainly in bioinformatics, is a workflow management system for reproducible data analyses. Rather than offering separate command-line tool wrappers, it employs a domain-specific language integrating Python and shell scripting within YAML-like code files. This reduces complexity in parsing command-line tool invocations but requires prior tool knowledge, and each tool invocation must be re-implemented within every workflow, limiting reproducibility.

Given the breadth of workflow engines receiving community support, we focus Styx on developing general-purpose, multi-language tool interfaces compatible with generic workflow engines.

Boutiques (Glatard et al., 2018) is a framework for FAIR command-line data processing. Its JSON-based Command-line Descriptor Standard contains: (i) tool metadata, (ii) command-line structure, (iii) input argument specification, (iv) output data specification, (v) argument groups and conditionals, and (vi) container information. The format’s strengths include machine-readability, JSON universality, interoperability between workflow systems, and automatic validation through JSON schema. Crucially for domains with significant tool heterogeneity, Boutiques is descriptive rather than prescriptive, meaning adoption doesn’t require modifying the tools being described, unlike standards like BIDS Apps that sought to homogenize tool interfaces (Gorgolewski et al., 2017).

The Common Workflow Language (CWL; Chapman et al. (2016)) is another standard for describing command-line tools and workflows using a declarative YAML/JSON format. CWL supports containerization and cloud computing environments and includes both tool description and workflow orchestration capabilities.

Historically, tool descriptions for Boutiques, CWL, and Nipype have been handwritten, requiring significant manual effort and domain expertise. We selected Boutiques as Styx’s backbone standard given its language independence and adoption in the neuroimaging community. However, our approach can extend to other standards both as truth sources and compilation targets, preserving compatibility with CWL, Nipype/Pydra Interfaces, and similar efforts.

## Methods

### Complexity of neuroimaging command-line interfaces

Formal language theory provides a framework for describing CLI structure (Hopcroft, 2001). A grammar defines valid token sequences that can appear in a CLI command. We propose neuroimaging CLIs can be effectively modeled using regular grammars, which are simple yet powerful enough to capture most CLI patterns.

While the Boutiques grammar describes simple command-line interfaces, many neuroimaging tools require more expressive power. We created a Boutiques standard superset that is a regular language, adding three key features: (i) Hierarchy: Nested subcommands (ii) Alternation: Mutually exclusive subcommands (iii) Repetition: Repeated subcommands. These features can be achieved by nesting Boutiques-like JSON objects within tool descriptors, preserving forwards-compatibility of existing Boutiques descriptors.

These extensions maintain Boutiques’ simplicity while providing necessary expressiveness. Figure 1 illustrates these concepts with examples from common neuroimaging tools. For the example shown in Figure 1d, which combines all three grammar features, we can express it using formal notation.

**Fig. 1.**
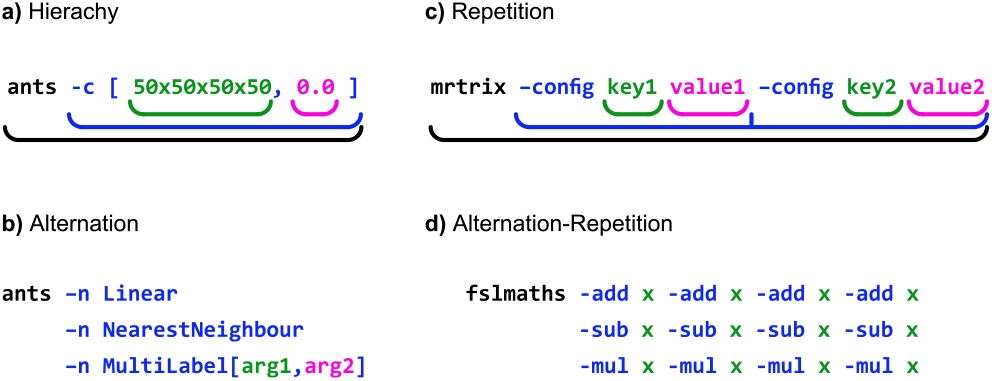
Examples of regular grammar features in neuroimaging CLIs. a) Hierarchy: ANTs N4BiasCorrection tool using nested arguments. b) Alternation: ANTs Registration tool’s interpolation argument. c) Repetition: MRTrix3 tools allowing arbitrary key-value pairs. d) Combined Alternation and Repetition: FSL’s fslmaths tool allowing a series of operations.

In the notation below, uppercase letters (A, B, C…) represent non-terminal symbols, angle brackets denote required parameters, the arrow *→* indicates production rules, *ε* represents an empty production (making elements optional), and the vertical bar | separates alternatives:

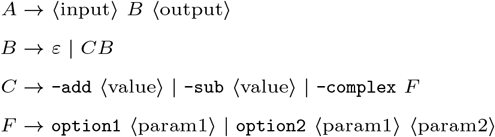

This formal representation captures how the tool accepts an input file, followed by zero or more operations (each with their own parameters), and finally an output file. The recursion in rule B allows for repetition of operations, while rule C shows alternation between different operation types, and F demonstrates hierarchical nesting of parameters.

By extending the Boutiques descriptor standard with formal language theory concepts, specifically regular grammars, we’ve developed a more expressive way to capture interface complexity. This approach provides a robust foundation for describing even the most complex neuroimaging CLIs. While the repetition extension is strictly necessary, the other implementations are theoretically possible without extensions but would significantly compromise code ergonomics.

The ability to express complex CLIs in a standardized, machine-readable format facilitates automatic wrapper generation, promotes interoperability between neuroimaging tools and workflow systems, and enhances reproducibility through clear tool interface descriptions.

### NiWrap — neuroimaging tool descriptions

Following the Boutiques standard extension, we created NiWrap as an organized collection of described CLI tools, defining criteria for tool selection and developing descriptor generation approaches.

The selection targeted all packages used by the neuroimaging pipelines C-PAC and fMRIPrep (Esteban et al., 2019), supplemented by tools for diffusion-weighted imaging (DWI) analysis, including Connectome Workbench and MRTrix3. Future inclusion of other packages is possible based on community demand.

Given the heterogeneity in neuroimaging CLI tools, no single approach was most efficient for creating tool descriptions. Figure 2 shows the paths taken for various tools. We employed two primary approaches: source code extraction and large language model generation. For Connectome Workbench and MRTrix3, we used source code extraction to programmatically analyze their source code and generate accurate descriptors. For remaining packages, we applied a hybrid approach combining Large Language Models (LLMs) and manual expert curation. Following these approaches, we generated nearly 2,000 command-line descriptions, summarized in Table 2.

**Table 2.** API coverage of generated wrappers across packages. These coverage metrics reflect descriptor availability rather than validated correctness. Ongoing efforts focus on refining and validating these descriptors, particularly those generated through the LLM-assisted approach.

| Package | Version | API Coverage |
| --- | --- | --- |
| AFNI | 24.2.06 | 565/611 (92.5%) |
| ANTs | 2.5.3 | 71/113 (62.8%) |
| Connectome Workbench | 1.5.0 | 202/202 (100%) |
| Convert3D | 1.1.0 | 2/2 (100%) |
| FSL | 6.0.5 | 239/310 (77.1%) |
| FreeSurfer | 7.4.1 | 707/800 (88.4%) |
| Greedy | 1.0.1 | 1/1 (100%) |
| MRTrix3 | 3.0.4 | 115/121 (95.0%) |
| MRTrix3Tissue | 5.2.8 | 1/1 (100%) |
| NiftyReg | 1.4.0 | 7/7 (100%) |
| dcm2niix | 1.0.20240202 | 1/1 (100%) |

**Fig. 2.**
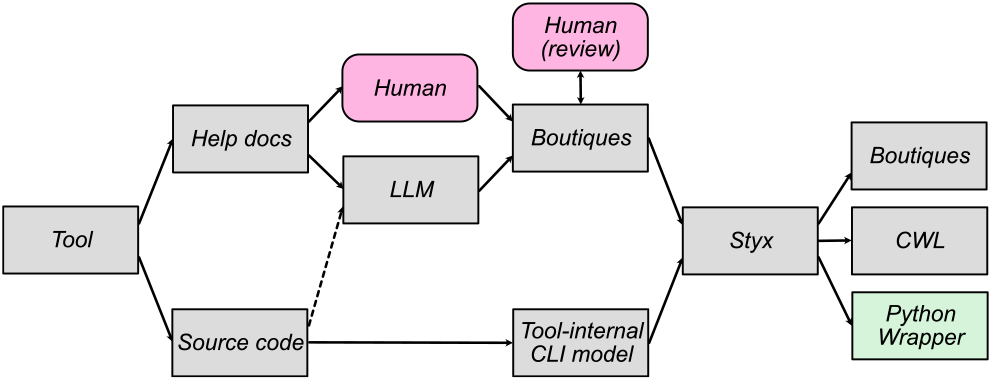
Approaches for large-scale Boutiques descriptor generation. The most direct path is extracting tool-internal CLI model data from source code. When impossible, tool help text was used (by humans or LLMs) to generate descriptors, requiring human review for correctness. As LLM context sizes grow, feeding entire tool source code to LLMs might increase accuracy.

#### Code extraction

When screening package source code, we discovered both Connectome Workbench and MRTrix3 implement abstractions around command line parsing suitable for direct extraction of data needed for Boutiques descriptors. With minor modifications to the source code, we leveraged these tools’ command-line interfaces to output structured metadata when passing the ‘– help’ flag. We then converted this JSON data to Boutiques descriptors. This process ensures high fidelity between tool implementation and descriptor, provides a sustainable update mechanism, and minimizes human error.

However, this method only applies to packages using centralized command-line parsing abstractions. Many neuroimaging packages, notably AFNI and FSL, hard-code command-line parsing individually for each tool, necessitating alternative approaches.

#### Large language models

For packages where direct source code extraction was infeasible, we employed large language models (LLMs: OpenAI GPT 3.5 and Claude 3.5 Sonnet) to bootstrap descriptor creation. We used single-shot prompting, providing an example pair of command-line help text and its corresponding Boutiques descriptor. We then prompted the model to generate draft descriptors for remaining programs by analyzing their help texts. After generation, we implemented automated post-processing of LLM outputs to ensure JSON structure validity.

### Styx Architecture

Given a collection of tool descriptions, Styx provides a native in-language API for consistent interaction across tools. We built the Styx compiler to create arbitrary language representations of each tool’s description.

#### Compiler Architecture

Styx employs a traditional compiler architecture with distinct frontend, intermediate representation (IR), and backend phases (Figure 3). This enables flexibility in both input formats and output languages while maintaining separation of concerns.

**Fig. 3.**
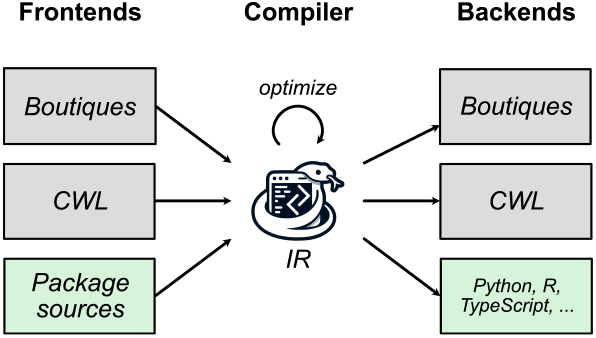
Architectural overview of the Styx compiler. The compiler follows a three-phase design: frontends, an intermediate representation (IR), and backends. Frontends (left) parse input from multiple sources: Boutiques descriptors (gray), Common Workflow Language (CWL) specifications (gray), and direct package source parsing (green). The compiler core maintains an IR that can be optimized through transformations (circular arrow). Backends (right) generate output in multiple formats: original descriptor formats like Boutiques and CWL (gray), and programming language bindings including Python, R, and TypeScript (green).

The frontend phase supports multiple input formats: (i) Boutiques descriptors (primary input), (ii) Common Workflow Language (CWL) tool specifications^1^, providing compatibility with existing bioinformatics community tool descriptions, and (iii) direct package source parsing for automated CLI specification extraction from well-structured codebases. Each frontend converts its input format into Styx’s internal representation.

The intermediate representation (IR) serves as a unified command-line interface model, capturing input parameters, output specifications, command-line construction rules, and documentation. This layer includes optimization that performs transformations like simplifying command patterns and normalizing parameter configurations. The IR is language-agnostic yet expressive enough to represent the full complexity of neuroimaging tool interfaces.

The backend phase generates code in target languages. Current implementations support Python, R, and TypeScript, with a modular design facilitating new target languages. Each backend implements translation rules converting the IR into idiomatic code, including type mapping from Boutiques/IR types to native language types, parameter validation logic, documentation strings, command-line argument formatting, and output file path resolution.

The backend can also generate Boutiques or CWL descriptions from the IR, enabling round-trip conversion between tool description formats.

#### Wrapper Integration Architecture

A key Styx feature is its decoupled execution model (Figure 4), separating command-line argument generation from tool execution. The architecture consists of three distinct layers:

**Fig. 4.**
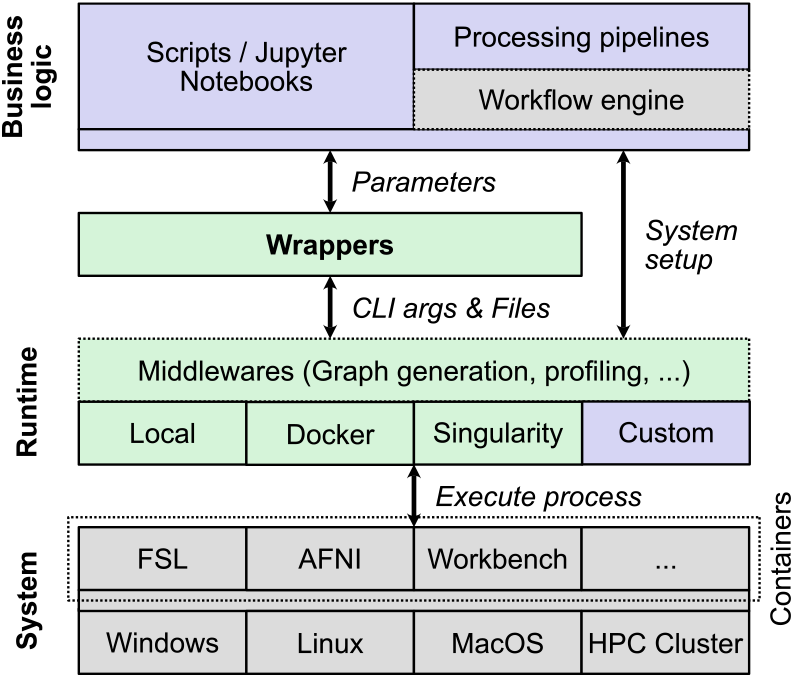
Systems diagram of Styx wrappers showing the layered architecture. Organized into three functional layers: Business logic (top, purple), Runtime (middle, light green), and System (bottom, gray). User scripts and pipelines interact with Styx wrappers, which communicate with the underlying system through middleware and various runtime environments. Green components represent Styx’s contributions, while gray components indicate third-party or system elements.

The business logic layer is where users write scripts, Jupyter notebooks, or processing pipelines, optionally using workflow engines. This layer passes parameters to wrappers and receives results.

The wrapper layer contains Styx-generated wrappers providing a consistent interface between user code and underlying systems, converting high-level function calls with named parameters into properly formatted command-line arguments and file specifications.

The minimal-overhead runtime layer handles actual execution across different environments, including middleware components for execution graph generation and resource profiling; runtime adapters for local execution, Docker/Singularity containers, or custom execution environments; system-level execution across operating systems or computing environments; and container-based isolation of neuroimaging tools.

This layered design enhances portability, allowing code written with Styx to run on any system regardless of native tool support. Execution is handled by interchangeable ‘runners’ tailored to specific environments. The middleware layer allows adding capabilities like execution tracking without modifying wrappers or user business logic.

The Styx compiler architecture provides a sustainable interface generation approach, centralizing the logic for creating in-language wrappers from Boutiques descriptors. This allows efficient adaptation to future target languages or command generation improvements, requiring modifications only to the compiler. A significant feature is the encoding of static information from Boutiques descriptors, including parameter and output names, types, and docstrings. This facilitates enhanced IDE functionality such as static analysis, auto-completion, and IntelliSense, improving code discoverability and early issue detection.

When used with container orchestration runners, Styx enhances reproducibility through sandboxing, ensuring commands can only access explicitly defined input files (read-only) and write to specified outputs. This prevents unintended file access or input modifications, providing guaranteed provenance.

Styx does not address workflow construction, but focuses on the definition of type-safe and reproducible interfaces. In cases where complex workflows are desired and needed, they can be seamlessly implemented atop Styx-compiled interfaces. This ability of Styx-compiled interfaces to be executed independent of workflow orchestration dramatically increases their portability and the range of possible use cases.

## Results

The primary output consists of a compiler and runtime system for generating and executing command-line tool wrappers, along with neuroimaging tool descriptors. The Styx compiler (https://github.com/styx-api/styx) generates programming language bindings from Boutiques descriptors, supporting Python, R, and JavaScript/TypeScript. Three runtime implementations are available: (1) local execution with shared type definitions (styxdefs), (2) Docker container runtime (styxdocker), and (3) Singularity container runtime (styxsingularity). When chaining wrapper calls, Styxgraph generates visualizations of file dependency graphs between operations. All components are MIT-licensed and available at https://github.com/styx-api. Comprehensive documentation and tutorials are provided at https://github.com/styx-api/styxbook.

The secondary contribution, NiWrap (https://github.com/styx-api/niwrap), provides over 1900 command-line descriptors covering 8 major neuroimaging packages, as detailed in Table 2.

### Structure of generated interfaces

The generated wrappers provide consistent, type-safe interfaces across programming languages. Below are examples of FSL’s BET being called through wrappers in different languages:

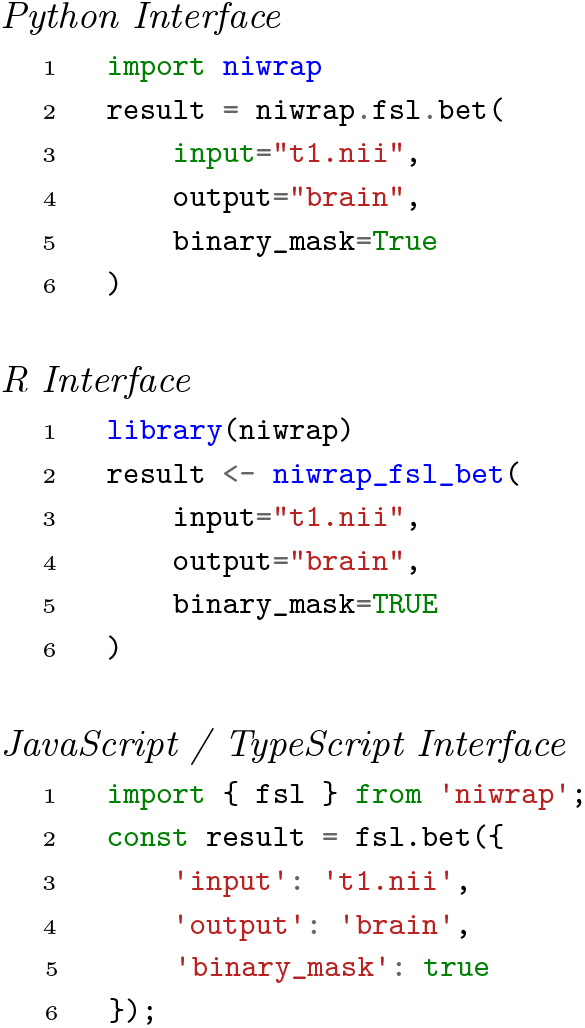

The wrappers implement static type information derived from Boutiques descriptors. Parameter types, default values, and documentation strings are encoded in function signatures, enabling IDE features like auto-completion. Return values are implemented as named tuples in Python, or equivalent structures in other formats, containing output file paths specified in the descriptor.

The compiler performs static analysis during wrapper generation to ensure type safety, including parameter type validation, required argument checking, and value constraint verification. Generated code is compatible with static type checkers such as MyPy, TypeScript’s type system, and R type-checking packages.

The runtime system implements a modular execution interface supporting local execution, Docker containers, and Singularity/Apptainer environments. Container configuration is handled through a standardized interface, with container specifications derived from tool descriptors.

### Evaluation of descriptor generation methods

Our approach to descriptor generation yielded different success rates depending on the source package. For MRTrix3 and Connectome Workbench, which implement centralized command-line parsing, we achieved almost complete API coverage through automated extraction—this approach can be easily repeated for new software versions, producing highly accurate descriptors reflecting tool implementation.

For packages without centralized parsing (AFNI, FSL, etc.), our LLM-assisted approach successfully generated initial descriptors for 60-90% of tools. No descriptors could be generated when help texts were missing or uninformative. Human review of LLM-generated descriptors is ongoing.

### Case study: pipeline development comparison

We present a case study comparing diffusion MRI preprocessing pipeline building using four approaches: Bash scripts, Nipype, Snakemake, and Styx with NiWrap. Each implementation performs standard preprocessing for diffusion MRI data with 10-15 command-line tools from packages like MRTrix3, FSL, and ANTs.

The implementations compared are:

- A Bash script-based pipeline (prepdwi) (Khan et al., 2021)
- A Nipype (version 1.0.3) workflow implementation (mrtpipelines) (Kai and Khan, 2022)
- A Snakemake-based (version 7.25.0) workflow (snakedwi) (Khan et al., 2023)
- A Styx & NiWrap (version 0.5.1) implementation (nhp-dwiproc) (Kai, 2025)

We focus on four components that highlight the strengths and weaknesses of each approach:

#### Data sourcing & syncing

File system I/O is necessary for all data processing pipelines, often requiring nuanced understanding of filenames and their relationships. All tools lack a uniform method for ingesting and outputting files, relying on string-substitution techniques. For Nipype, Snakemake, and Styx, external packages such as bids2table or pybids can standardize file system I/O.

#### Pipeline branching

In Styx and Bash, conditional branching follows standard programming paradigms (e.g., if −else statements), allowing familiar control flow structures.

In Nipype, workflows consist of Nodes associated with each interface. Implementing conditional branching requires dividing workflows into smaller sub-workflows at each decision point, adding structural complexity as conditions increase.

For Snakemake, nodes connect implicitly via matching input and output file names. Two nodes connect when the output file of one matches the input file name of another. This approach provides flexibility for branching points while following conditional paradigms.

#### Workflow Management

Bash and Styx don’t provide built-in workflow engines, so caching and parallelization aren’t native features. Both frameworks support integration with external tools, allowing developers to select appropriate workflow engines.

Nipype and Snakemake incorporate workflow engines with integrated caching and parallelism. Both check for existing output files before executing a workflow and skip interfaces when outputs are present. Nipype implements parallelism through `MapNodes` and `iterables`. `MapNodes` enable parallelization across a single interface, while `iterables` facilitate parallelization across entire workflows. Snakemake offers similar parallelization through wildcard substitutions and iterable data types.

#### Debugability

Common pipeline construction challenges include resolving individual interface errors and addressing misconnections between workflow nodes. Interface errors typically manifest as crashes with error codes and messages. All frameworks allow displaying these errors in the terminal or directing them to log files, making it possible to trace specific commands that lead to failures.

More challenging are errors from incorrect connections between components or skipped nodes. In Bash, these issues often stem from incorrect variable passing, requiring manual identification of workflow failure points. Styx requires user identification of missed connections, though its middleware supports graph visualization to help locate connection problems. Nipype’s explicit connections with graph visualization facilitate identifying missed connections. Snakemake can generate graphs to assist with diagnosing connectivity issues, but its implicit file-based connections can complicate troubleshooting when connections fail.

Styx provides pre-runtime validation through type-checking and static code analysis, helping identify issues like missing upstream files or variables before execution. While other frameworks technically support validation, their reliance on string-formatted data limits these features’ effectiveness.

Each framework presents different learning curves. Bash requires shell scripting familiarity. The Python-based tools require Python knowledge, but Nipype and Snakemake demand additional understanding of their workflow engines and execution plugins. Snakemake further uses configuration files in JSON/YAML and employs a Python-based domain-specific language. Styx, like Bash, relies primarily on base language knowledge without requiring familiarity with additional domain-specific languages or workflow architectures—a significant advantage, though with the trade-off of having no embedded workflow engine.

## Discussion

The coverage of 62% − 100% across major neuroimaging packages demonstrates our approach’s viability for CLI wrapper generation. Particularly significant is the complete (95% − 100%) coverage of MRTrix3 and Connectome Workbench through direct source code extraction. This automated approach ensures accuracy and provides a sustainable path for maintaining wrappers as tools evolve: when these packages release updates, new descriptors can be automatically generated from their source code. For packages where direct extraction wasn’t possible, our LLM-assisted approach proved effective, though requiring manual verification. This hybrid approach demands more maintenance effort than direct extraction but significantly improves upon fully manual descriptor creation. The compiler-based architecture means structural improvements or new features can propagate to all wrapped tools across all supported languages through changes to a single codebase, providing a highly maintainable solution.

The regular grammar extension to Boutiques successfully captured neuroimaging CLI complexity, including hierarchical structures, alternating options, and repeated arguments. This comprehensive modeling allows Styx to generate wrappers that preserve full tool functionality while presenting them through a more accessible interface. Complex tool invocations that might require multiple shell script lines can now be expressed as single, type-safe function calls with clear parameter documentation.

A key achievement is the seamless integration of existing command-line tools into modern Python-based workflows. The generated wrappers enable pipeline development in pure Python without shell scripting, while leveraging modern features such as IDE auto-completion and type checking. This integration bridges the gap between traditional neuroimaging software and contemporary data science practices, potentially accelerating new analysis pipeline development.

While Styx successfully addresses many CLI tool integration challenges, several limitations should be noted. The wrappers can be used with any Python-based parallel processing solution but don’t provide built-in distributed computing capabilities. While this is a limitation relative to similar solutions, we believe this decision increases Styx’s portability and long-term usability. Several opportunities for extension and improvement exist. Currently generating wrappers for Python, R, and JavaScript/TypeScript, the latter’s static typing system aligns naturally with our approach and could provide strong type safety guarantees for neuroimaging pipelines in web-based environments. This complements the growing ecosystem of browser-based visualization and analysis tools, though these wrappers would still require a backend server to execute the actual command-line tools.

Adding content type information to file inputs and outputs would enable tighter integration with modern neuroimaging libraries, potentially allowing direct interfacing between command-line tools and in-memory data structures. This could be implemented using ramdisks or shared memory segments, enabling tools to operate on data without requiring disk I/O between processing steps, significantly improving performance for complex pipelines with many sequential operations.

Automated testing of generated wrappers presents challenges: without content type information, determining appropriate test inputs is difficult; large parameter spaces make exhaustive testing impractical; and long runtimes combined with ambiguous error reporting make it difficult to differentiate between implementation errors and invalid parameter combinations.

These challenges may require tool creators’ involvement to address efficiently. As the project matures, broader adoption would naturally lead to improved tool coverage and descriptor accuracy through real-world usage and feedback.

Beyond neuroimaging, the approach demonstrated by Styx and NiWrap could adapt to other scientific domains facing similar challenges with legacy command-line tools, such as those in bioinformatics. The combination of formal grammar-based interface description and automated wrapper generation serves as a template for modernizing scientific software ecosystems while preserving valuable legacy implementations.

Several future development directions are planned for Styx. First, completing the CWL frontend and backend implementations will expand interoperability with the bioinformatics community where CWL has significant adoption. This would allow seamless translation between Boutiques and CWL descriptions, creating a bridge between neuroimaging and bioinformatics tool ecosystems. Second, we plan to develop content-aware interfaces that understand the semantic meaning of files, potentially allowing direct memory-based data exchange between tools rather than requiring intermediate file I/O. Finally, we aim to implement automated validation pipelines for generated wrappers to ensure both syntactic and functional correctness across the full range of supported tool interfaces.

## Conclusion

We present Styx, a compiler architecture to create in-language interfaces for command-line tools. This approach involves describing tools using a Boutiques tool description standard superset, parsing these descriptions into an intermediate representation, before compiling them into an extensible list of backends. Styx minimizes the complexity of adopting command-line tools within commonly-used analytic programming environments and adds significant standardization across interfaces of tools from distinct packages with unique nomenclatures and command-line structures. NiWrap, our collection of over 1900 descriptors, demonstrates the scalability of using both source code modification and LLM-assisted generation to create valid tool descriptions and lowers the barrier for tool integration within the scientific software ecosystem. Together, Styx and NiWrap offer a sustainable solution for bridging legacy command-line interfaces with modern programming paradigms, enabling researchers to focus more on scientific questions and less on technical implementation details.

## Data and Code Availability

All software described in this manuscript is open source and available under the MIT license:

- Styx compiler and runtime: https://github.com/styx-api/styx
- NiWrap tool descriptors: https://github.com/styx-api/niwrap
- Documentation and tutorials: https://github.com/styx-api/styxbook

The case study pipelines compared in this work are available at:

- prepdwi (Bash): https://zenodo.org/records/4959832
- mrtpipelines(Nipype): https://github.com/kaitj/mrtpipelines
- snakedwi(Snakemake): https://zenodo.org/records/8237288
- nhp-dwiproc (Styx): https://github.com/HumanBrainED/nhp-dwiproc

## Competing interests

No competing interest is declared.

## Author contributions statement

F.R. conceived and implemented the Styx compiler system, the architecture, and developed the core functionality. J.K. and B.S. contributed to the development of tool descriptors for NiWrap. J.K. assisted in manuscript preparation. T.X., S.G., T.G., M.P.M., and G.K. provided senior oversight and guidance throughout the project. G.K. contributed substantially to manuscript preparation and editing. All authors reviewed and approved the final manuscript.

## Acknowledgments

Research reported in this publication was supported in part by the National Institute of Mental Health under Award Number RF1MH130859 (G.K., M.P.M.). The content is solely the responsibility of the authors and does not necessarily represent the official views of the National Institutes of Health.

We thank Connor Lane for helpful feedback and discussions during the development of this project.

## Footnotes

1 The CWL frontend, and backend are still a work in progress.

## Notes

https://github.com/styx-api

### Competing Interest Statement

The authors have declared no competing interest.

